# The maturity in fetal pigs using a multi-fluid metabolomic approach

**DOI:** 10.1101/2020.03.13.990564

**Authors:** Gaëlle Lefort, Rémi Servien, Hélène Quesnel, Yvon Billon, Laurianne Canario, Nathalie Iannuccelli, Cécile Canlet, Alain Paris, Nathalie Vialaneix, Laurence Liaubet

**Author notes:** these authors contributed equally to this work (Nathalie Vialaneix, Laurence Liaubet).

## Abstract

In mammalian species, the first days after birth are an important period for survival and the rates of mortality before weaning are high. In pigs, the perinatal deaths average 20% of the litter, with important economic and societal consequences. Among the factors influencing piglet survival at birth, the maturity is likely to be one of the most important. Maturity can be defined as the outcome of complex mechanisms of intra-uterine development and maturation occurring during the last month of gestation. Here, we provide new insights on maturity by studying the end of gestation at two different stages (three weeks before term and close to term) in two breeds of pigs that strongly differ in terms of neonatal survival. Since metabolomics is a promising approach for phenotype characterization or biomarker discovery, we provide a complete understanding of the metabolome of the fetuses in late gestation in three fluids (plasma, urine, and amniotic fluid). We found that biological processes related to amino acid and carbohydrate metabolisms are critical for piglet maturity. We also confirmed some previously described metabolites associated with delayed growth (*e.g.*, proline and myo-inositol). Altogether, our study proposes new routes for a better characterization of piglet maturity at birth.

## Introduction

In mammalian species, the first days after birth are an important period for survival and the rates of mortality before weaning are high. In humans, even if the rate of mortality has strongly decreased in the recent years, neonatal deaths (before one month after birth) still represents 47% of the death before five years old (about 2.5 million per year) [53]. In a polytocous species like swine, this rate averages 20% of the litter [16], with the most critical time for piglet survival during the perinatal period that includes the first 24 hours of life. In pigs, a substantial increase in piglet mortality before weaning can be attributed to the effect of animal selection: since improving the prolificacy rate and the carcass merit (*i.e.*, leaner meat) has been the main objectives of selection, the undesired effect may be a loss of robustness in animals, a greater susceptibility to disease, a weakened locomotion and behavioural problems [44, 46]. From a physiological point of view, many factors have been identified as influencing piglet survival [16]. They have been related to maternal effects (*e.g.*, intrauterine effects, farrowing duration, parity, health status), to piglet factors (*e.g.*, genetic type, vitality at birth) or even to piglet characteristics that are partly under maternal control (*e.g.*, birth weight). Among those factors, piglet maturity is likely to be an important determinant of subsequent survival and postnatal growth [2, 31], which has also been proved altered by animal selection [10]. Maturity at birth, which can be defined as the complete development allowing survival at birth, is the outcome of complex mechanisms of intra-uterine development and maturation occurring during the end of gestation [31]. In pigs, the maturation is known to occur during the last month of gestation (approximately 90-114 days of gestation, dg). Physiological maturity at birth together with environmental conditions thus have important consequences on neonatal mortality. More precisely, Bauer *et al.* [3] have reported that IntraUterine Growth Restriction (IUGR) is responsible for part of neonatal mortality and reduced growth later in life in a mixed German domestic breed. IUGR is thus directly associated with maturity at birth and it is not surprising that birth weight is often considered as an important indicator for piglet perinatal survival. However, this relation between birth weight and maturity at birth was observed within a given genotype but not accross genotypes: for instance, piglets from the Chinese Meishan (MS) breed exhibit a greater maturity at birth and a better rate of survival than piglets from the Large White (LW) breed, which have larger birth weights. This highlights the need for a deeper understanding of maturity to effectively reduce perinatal moratlity.

In this context, the end of gestation was previously studied in two breeds of pigs that strongly differ in terms of neonatal survival [56]. The LW breed, which represents the European breeds and has been genetically selected for lean growth and prolificacy, presents a high rate of neonatal mortality partly due to a lower physiological maturity at birth. On the contrary, the MS breed presents a low rate of mortality and was considered as more mature at birth [9, 11]. LW and MS sows were inseminated with a mixed semen (LW and MS) so that pure and crossed fetuses grew in the same uterine environment. The fetuses were studied at two key gestational stages, 90 days being the onset of fetal maturation for muscle [20] and 110 days being close to term. Samples collected corresponded to eight different conditions (four genotypes – LW, MS, LW×MS, and MS×LW – and two gestational stages). Samples collected at 110 dg are thus representative of an increased maturity compared to 90 dg, as are data collected on MS compared to LW. Crossed fetuses were expected to present intermediate maturity between LW and MS fetuses, in general, as already shown in the previous transcriptome study [56]. Maternal or paternal effects or heterosis led to some exceptions for certain metabolites. These exceptions will be systematically discussed.

The transcriptomic and/or proteomic analyses of the muscle, the intestine and subcutaneous adipose tissue underlined a switch in gene expressions between the two stages of gestation and a delay in the development of tissues and organs in LW piglets. Notably, genes involved in muscular development are up-regulated around 90 dg and genes linked to metabolic functions, like gluconeogenesis, are up-regulated at 110 dg either in muscle or adipose tissue whatever the genotype [22, 55, 56, 66].

The present study uses the same experimental protocol and aims at completing the transcriptomic and proteomic analyses by providing a complete understanding of the metabolome of the fetuses in late gestation. Metabolomic is a promising approach to investigate health and welfare in large cohorts, for phenotype characterization or biomarker discovery. Indeed, high-throughput metabolome measurements can be obtained easily and at an affordable cost by ^1^H Nuclear Magnetic Resonance (NMR) and the metabolome provides a comprehensive characterization of the small molecules involved in metabolic chemical reactions. It is thus an interesting approach to identify usable biomarkers. Although some blood parameters have already been associated with piglet maturity at birth (*e.g.*, albumin and IGF-I plasma concentrations [10]), a comprehensive study was needed to explore the metabolic pathways involved in prenatal maturation and to identify new biomarkers of maturity that may be used at birth. To achieve this aim, we compared the metabolomes of three fluids, namely plasma, urine and amniotic liquid, of more than 600 fetuses obtained at 90 or 110 dg. These three fluids were chosen to represent different aspects of the fetus metabolism: the plasma reflects the metabolic regulations of the fetus while the urine metabolome reflects its excretory renal function and the amniotic fluid reflects its nutritional function and mechanic protection as well as the interaction with maternal and placenta tissues.

To measure metabolite concentrations in the three fluids, NMR metabolomic spectra were treated with **ASICS**, a recently developed R package [32]. Discriminant analyses and mixed models were applied to identify potential biomarkers of maturity at birth in the different fluids. Systematic enrichment analyses were performed on KEGG pathways to identify those that were the most involved in fetal development during late gestation. This allowed us to confirm some previously described metabolites associated with delayed growth and to identify important biological processes involved in piglet maturity.

## Results

Among the 190 metabolites available in the reference library of **ASICS** package, about 65 metabolites were identified for each fluid (*i.e.*, 63 for plasma, 64 for urine and 68 for amniotic fluid; Supplementary Table S1 and Supplementary Fig. S1). Thirty-nine metabolites were identified in all fluids including many amino acids (*e.g.*, glutamine, glycine, proline and arginine) and many sugars (*e.g.*, glucose, fructose, glucose-6-phosphate). Other metabolites were identified in only one or two fluids like leucine and isoleucine (that were identified only in plasma and urine) or reduced or oxidated glutathione (identified only in urine and amniotic fluid).

### Multivariate exploratory analyses

Three Orthogonal Projections to Latent Structures Discriminant Analyses (OPLS-DA)[52], one for each fluid, allowed discriminating the two stages of gestation with a good accuracy (Fig. 1 and Supplementary Fig. S2), especially in the plasma where the cross-validation error was equal to 1%. For urine and amniotic fluid, the error was a bit higher (4%) but still low, which indicated a slightly weaker separation between the two groups (90 dg and 110 dg). Altogether, these results suggested that OPLS-DA could be interpreted with a high level of confidence.

**Figure 1:**
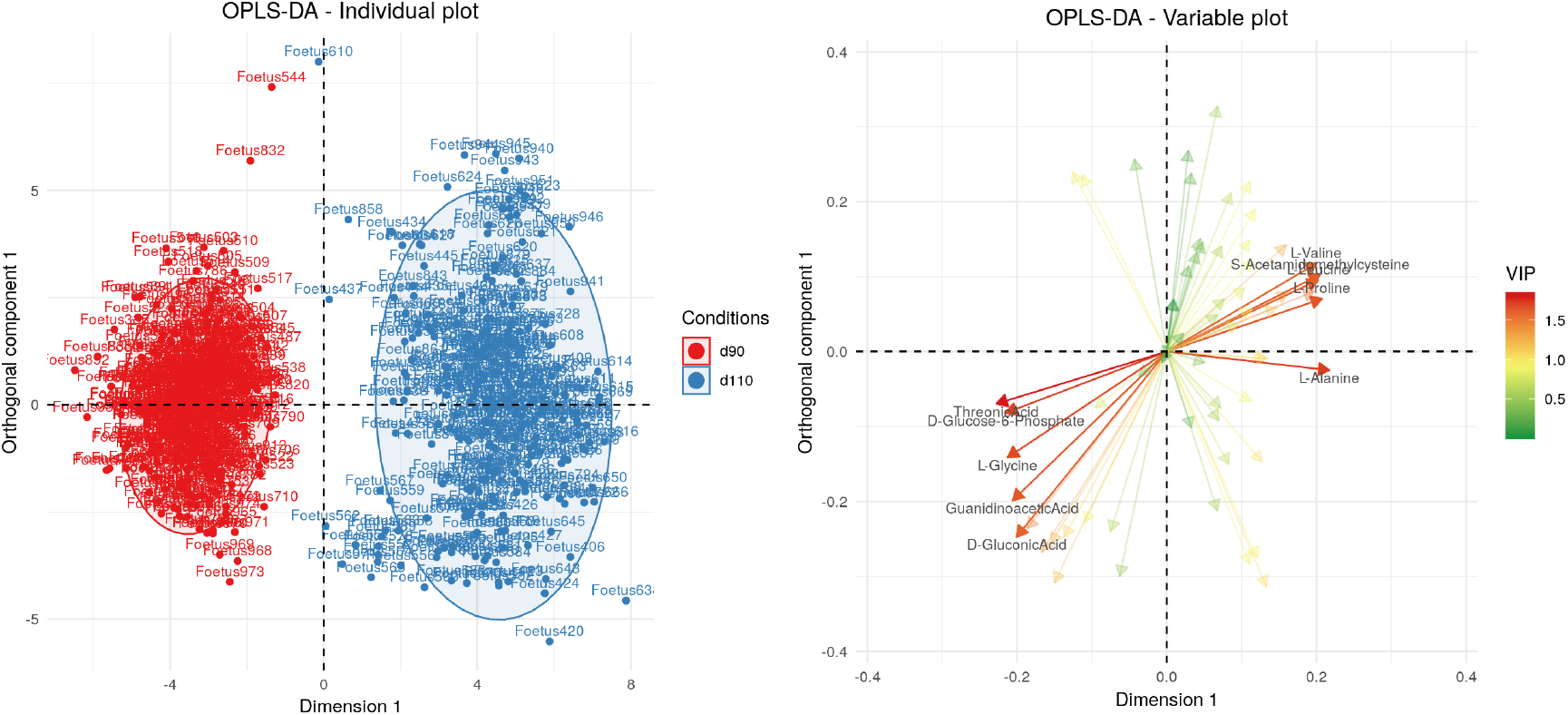
Individual and variable plots of the OPLS-DA on plasma spectra.

About twenty metabolites were found influential (Variable Influence on Projection, VIP ¿ 1) for each fluid (23 for plasma, 21 for urine and 22 for amniotic fluid) and used in enrichment analyses of pathways. Results of these analyses are presented in Table 1 and were performed for each fluid separately. Only three influential metabolites were common to the three fluids (glucose-6-phosphate, fructose and guanidinoacetate; Supplementary Fig. S3). Guanidinoacetate and fructose were more concentrated at 90 dg than at 110 dg in the three fluids while glucose-6-phosphate was more concentrated at 90 dg in plasma and amniotic fluid but more concentrated at 110 dg in urine. Glucose-6-phosphate is included in the “galactose metabolism”, a pathway that was found enriched in influential metabolites for all fluids and was also part of the “pentose phosphate pathway” which was enriched in influential metabolites in plasma. With the fructose, glucose-6-phosphate is also involved in the “starch (*i.e.*, glycogen in animals) and sucrose metabolism”. This pathway was enriched in influential metabolites in both urine and amniotic fluid. However, concentrations of all influential metabolites of this pathway varied differently in the two fluids between the two gestational stages. In urine, most of these metabolites had a higher concentration at 110 dg (glucose-6-phosphate, glucoronate and maltose) but in amniotic fluid, only the glycogen has a higher concentration at 110 dg.

**Table 1:**
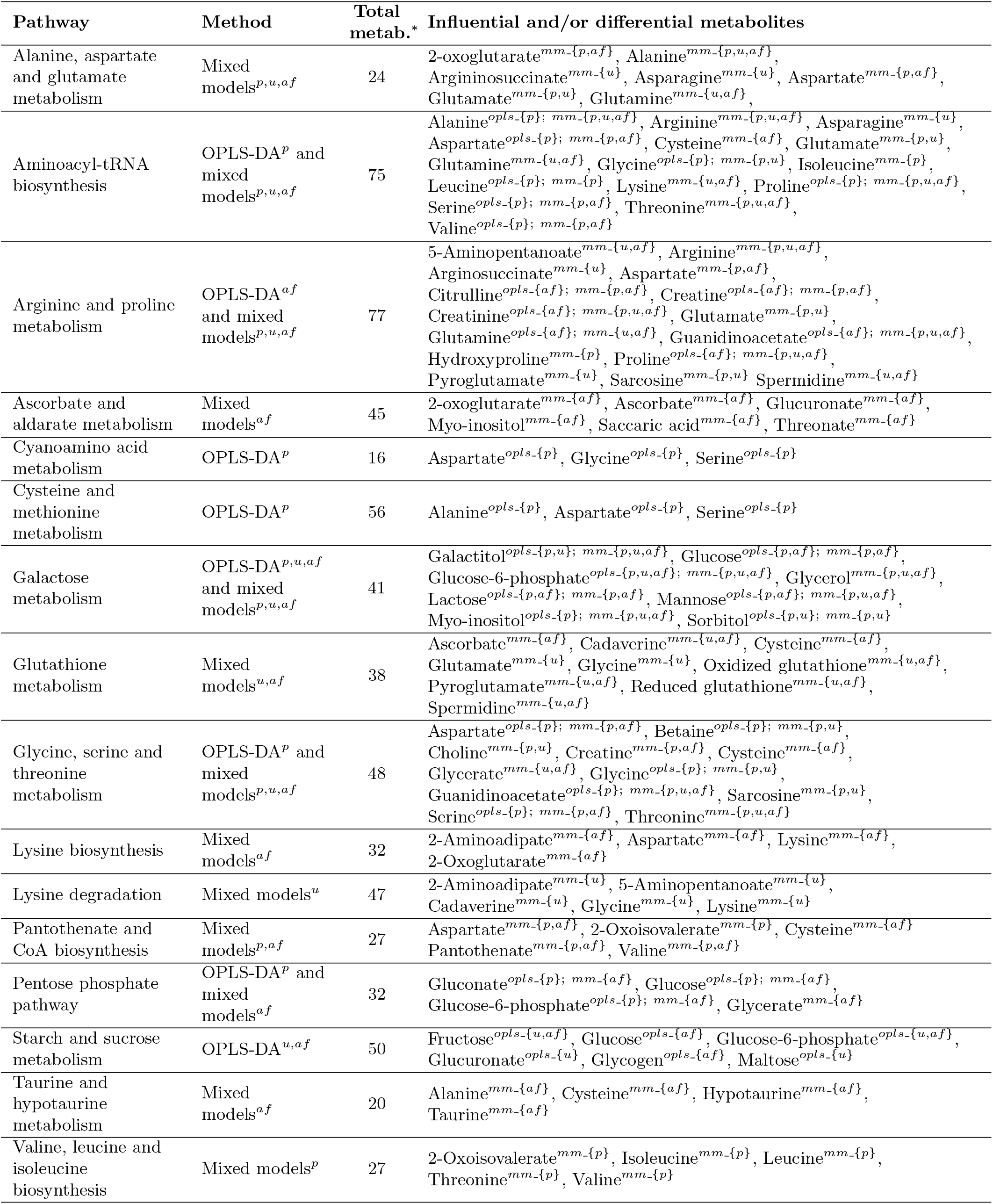
Enriched pathways in influential metabolites for OPLS-DA and in differential metabolites for mixed models. ^<*method*>−{<*fluids*>}^ influential and/or differential metabolite for <method> (^*opls*^ for OPLS-DA and ^*mm*^ for mixed models) in <fluids> (*p* for plasma, *u* for urine and *af* for amniotic fluid) * Total number of metabolites in the pathway.

Finally, guanidinoacetate is involved in the metabolic pathways of several amino acids. In addition, six amino acids were found influential only in plasma (alanine, aspartate, glycine, leucine, serine and valine). The pathway enrichment analysis highlighted six pathways enriched in influential metabolites in plasma, among which four were related to amino acids: “aminoacyl-tRNA biosynthesis”, “glycine, serine and threonine metabolism”, “cyanoamino acid metabolism” and “cysteine and methionine metabolism”. In amniotic fluid, a pathway related to amino acids was also found enriched in influential metabolites: the “arginine and proline metabolism”.

### Differential analyses

Mixed linear models were fitted to each metabolite independently. The complete model involved two factors, gestational stage and fetal genotype, as well as their interaction (fixed effects), and the sow as a random effect. In addition, all differential metabolites (*i.e.*, metabolites for which the complete model was significantly better than the model with just the sow effect) were submitted to a pathway enrichment analysis. They were then associated with one of the best-fit sub-model derived from the complete model to help their individual interpretation (see Methods). All the influential metabolites extracted by the multivariate analysis were also found significantly differential for one of these mixed linear models, whatever the fluid. Detailed results of mixed models are available in Supplementary Data 1.

Mixed models highlighted that metabolomes differed more between the two gestational stages than between genotypes, for all fluids. In plasma, 57 differential metabolites were found associated with a model that included a stage of gestation effect (complete, additive and only stage models) whereas only 28 differential metabolites were found associated with a model that included a genotype effect (complete, additive and only genotype models). In urine and amniotic fluid, the comparison respectively gave 41 versus 20 metabolites and 58 versus 6 metabolites (Table 2).

**Table 2:** Number of differential metabolites associated with each sub-models by fluid.

| Sub-model | Plasma | Urine | Amniotic fluid |
| --- | --- | --- | --- |
| Complete | 8 | 0 | 0 |
| Additive | 15 | 5 | 4 |
| Only stage | 34 | 36 | 54 |
| Only genotype | 5 | 15 | 2 |
| **Total for models with a stage effect** | **57** | **41** | **58** |
| **Total for models with a genotype effect** | **28** | **20** | **6** |
| **Total** | **62** | **56** | **60** |

#### Differences between stages of gestation

More differential metabolites were found associated with a model with a stage of gestation effect (complete, additive and only stage models) in plasma than in urine and amniotic fluid but, in amniotic fluid, 54 differential metabolites (out of 58) were associated with the model with only the stage effect, which is the highest number of differential metabolites for this model among the three fluids. In addition, over-time changes of quantifications for metabolites associated with the only stage model were different in plasma than in the other fluids. A majority of metabolites (28/34) were more concentrated at 110 dg than at 90 dg in plasma whereas only half of the metabolites were more concentrated at 110 dg than at 90 dg in urine and amniotic fluid.

On the 20 proteinogenic amino acids, 15 amino acids were found differential in at least one fluid. All differential amino acids were associated with a stage effect model, except for five: in urine, alanine, glutamine, glycine, proline and threonine were associated with the only genotype model. These 15 amino acids are included in four pathways that were found enriched in differential metabolites in the three fluids (Table 1), namely the “alanine, aspartate and glutamate metabolism”, the “aminoacyl-tRNA biosynthesis”, the “arginine and proline metabolism” and the “glycine, serine and threonine metabolism”. Metabolites in these pathways were more concentrated at 110 dg for 14 out of 20 metabolites in plasma and for 11 out of 18 metabolites in amniotic fluid.

However, differences were also identified between the fluids. In plasma, 2-oxoisovalerate, isoleucine, leucine, threonine and valine were all found differential and associated with the only stage model. They are included in the “valine, leucine and isoleucine biosynthesis”, a pathway related to amino acids that was found enriched in plasma. In amniotic fluid, 2-aminoadipate, aspartate, lysine, and 2-oxoglutarate were differential and all found associated with the only stage model. They are included in another pathway related to amino acid, the “lysine biosynthesis”, which was found enriched in amniotic fluid. Metabolites of these two pathways (“valine, leucine and isoleucine biosynthesis” and “lysine biosynthesis” pathways) were all more concentrated at 110 dg.

In urine, the number of differential metabolites associated with a model with a stage effect was smaller than in the other two fluids, especially for amino acids. However, the “galactose metabolism”, which was found enriched in differential metabolites for the urine, contained five metabolites (myo-inositol, glucose-6-phosphate, mannose, sorbitol and galactitol), which were all associated with the only stage model.

Finally, also in amniotic fluid, four differential metabolites were found associated with a model including a stage effect: glucose-6-phosphate, gluconate, glucose and glycerate, among which 3 out of 4 (except for the gluconate) were associated with the only stage model. These metabolites are all included in the “pentose phosphate pathway”, which was found enriched in differential metabolites in amniotic fluid. This pathway had also been previously found enriched in influential metabolites (as obtained by OPLS-DA) but in plasma rather than in amniotic fluid. In addition, two metabolites of this pathway, glucose and gluconate, varied in opposite directions in the two fluids: glucose concentration was higher at 110 dg in plasma whereas it was higher at 90 dg in amniotic fluid (and the variation was the opposite for gluconate concentration).

#### Differences between genotypes

More differential metabolites were found associated with a model that included a genotype effect (complete, additive and only genotype models) in plasma than in urine and amniotic fluid (28 metabolites compared to 20 and 6, respectively; Table 2).

In plasma, six differential metabolites (galactitol, glucose, glucose-6-phosphate, mannose, myoinositol and sorbitol) were found associated with the complete or the additive model, which included a genotype effect, and two (glycerol and lactose) with the only stage model. Among these 8 metabolites, some were also found differential for urine and amniotic fluid but they were usually not associated with a model including a genotype effect in these fluids. The only exceptions were the glycerol in urine (associated with the only genotype model) and the galactitol in amniotic fluid (also associated with the only genotype model). In addition, these 8 metabolites are all included in the “galactose metabolism” pathway, which was found enriched in differential metabolites in all fluids (Fig. 2). In plasma, mannose and glucose were more concentrated at 110 dg than at 90 dg and they were also more concentrated in MS than in LW at both 90 dg and 110 dg. On the contrary, the other three metabolites (glucose-6-phosphate, sorbitol and myo-inositol) were more concentrated at 90 dg and 110 dg in LW and the galactitol was more concentrated in MS at 110 dg. In addition, the concentration of myo-inositol was higher when the fetus had a LW father (whatever the mother genotype) and the concentration of glucose-6-phosphate, sorbitol and galactitol was higher when the fetus had a MS father. On the contrary, in urine, the glycerol had an higher concentration when the fetus had a LW mother.

**Figure 2:**
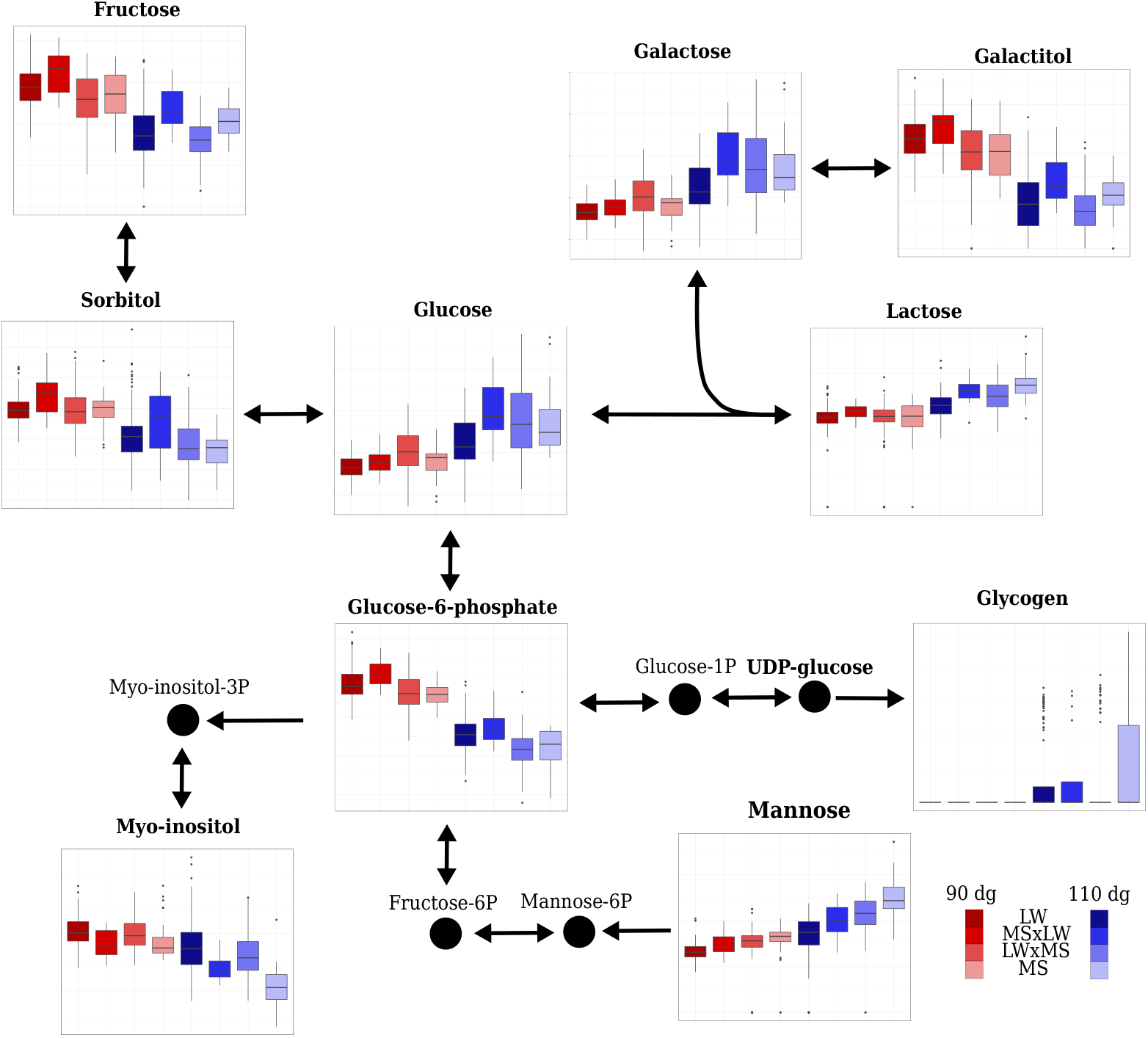
Plasma relative concentrations of some metabolites involved in the carbohydrate metabolism pathways (“galactose metabolism” and “starch and sucrose metabolism”) according to stage of gestation and fetal genotype. In bold, metabolites included in **ASICS** reference library. The coordinates of the *y* axes in boxplots can not be compared between two metabolites (relative concentrations limits of the boxplots are adapted to each metabolites).

In plasma and urine, 12 differential metabolites of the enriched “arginine and proline metabolism” and “glycine, serine and threonine metabolism” pathways were also found associated with a model including a genotype effect: the creatinine (complete model), the aspartate, the glycine, the guanidinoacetate and the proline (additive model), the choline and the creatine (only genotype model) in plasma and the glutamate and the guanidinoacetate (additive model), the 5-aminopentanoate, the glutamine, the glycerate, the glycine, the proline and the threonine (only genotype model) in urine (see Fig. 3 for a representation of creatine, creatinine, glutamate, glutamine, glycine, guanidinoacetate and proline). Among these metabolites, 6 had a higher concentration in MS than in LW (5-aminopentanoate and glycerate in urine, aspartate, choline and creatine in plasma and proline in both urine and plasma). Concentrations of aspartate in plasma and glycerate in urine were also higher when the fetus had a MS father compared to a LW father (paternal effect; Supplementary Fig. S4), while median concentrations for choline and creatine in plasma were larger than 0 only for fetuses with a MS mother and father (effect of the pure MS genotype). Three metabolites (glutamine in urine and glycine and guanidinoacetate in plasma and urine) had a higher concentration in LW than in MS.

**Figure 3:**
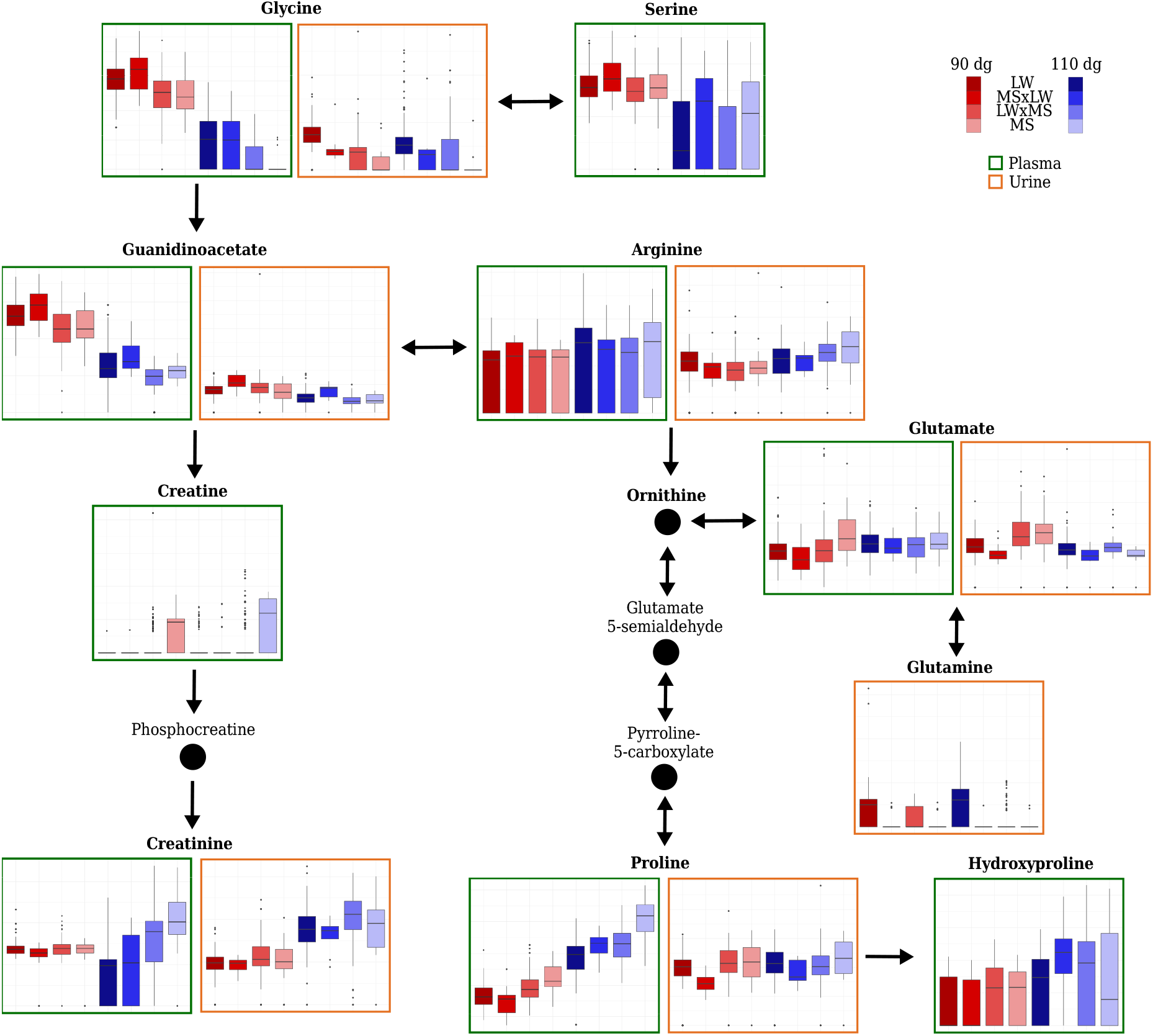
Plasma and urine relative concentrations of some metabolites involved in amimo acid metabolism (“arginine and proline metabolism” and “glycine, serine and threonine metabolism”) according to stage of gestation and fetal genotype. For the sake of clarity, only 9 and 7 metabolites are represented out of 15 found differential in plasma and 15 found differential in urine. In bold, metabolites included in **ASICS** reference library. The coordinates of the *y* axes in boxplots can only be compared between plasma and urine for the same metabolite but not between metabolites (relative concentrations limits of the boxplots are adapted to each metabolites).

In urine, four differential metabolites associated with the enriched “glutathione pathway” (Fig. 4) were associated with the only genotype model (the oxidized glutathione, the glycine, and the pyroglutamate) and one was associated with the additive model (the glutamate). These metabolites were more concentrated in LW than in MS fetuses at 110 dg and more concentrated in MS than in LW at 90 dg, except for the glycine (that remained more concentrated in LW than in MS at 90 dg).

**Figure 4:**
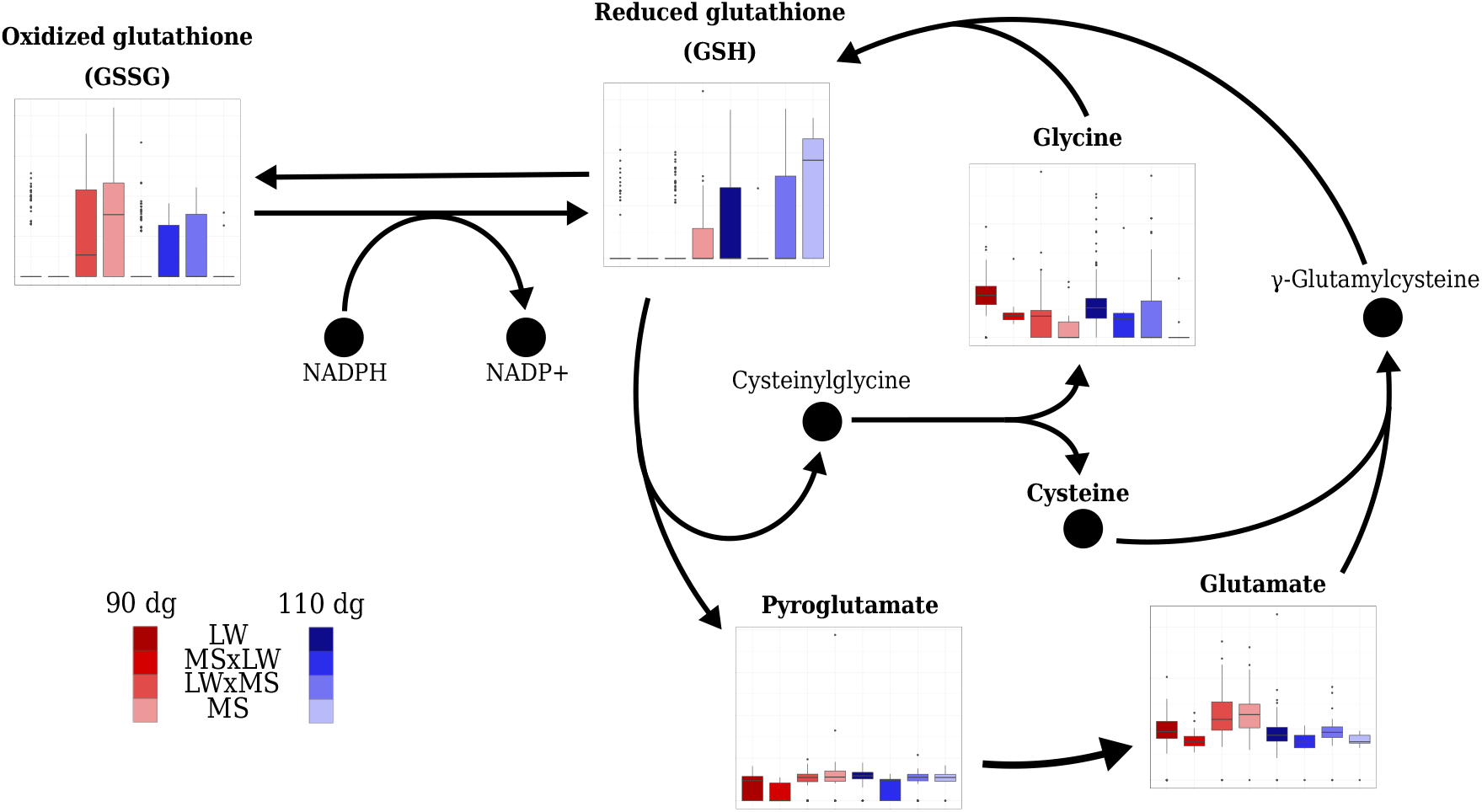
Urinary relative concentrations by stages of gestation and genotypes for some metabolites of “glutathione pathway” (for the sake of clarity, only the *γ*-glutamyl-cycle is represented in this figure). In bold, metabolites included in **ASICS** reference library. The coordinates of the *y* axes in boxplots can not be compared between two metabolites (relative concentrations limits of the boxplots are adapted to each metabolites).

## Discussion

Investigating the biological processes of fetal maturation and fetal growth retardation is of major interest in humans and also in several mammalian livestock species, like sheep[24] and pig[50, 57]. These characteristics are related to fetal development during late gestation, which is difficult to explore in mammalian species due to the invasiveness of experiments performed during that period. Hence, since impaired maturation may induce postnatal developmental delay, metabolic syndrome, or early death[23], the evaluation of the level of development at birth is currently done by measuring the birth weight[4] as a proxy for intrauterine development. The study of the metabolome in the late gestation is a promising approach to predict immediate or later outcomes and to evaluate fetal growth retardation[33].

In humans, some metabolomic studies have already been performed on amniotic fluid collected during the amniocentesis at the second or the third trimester of gestation[19, 25]. However, most of the other metabolomic surveys in humans have been performed later at birth, especially on cord blood[17, 18, 48], or on urine[14]. In non-human mammalian species, few studies related to the metabolome during late developmental processes have been published so far. Among them, a plasmatic NMR study has been performed on late pig fetuses[40].

Using NMR techniques, we acquired untargeted metabolomic measurements on three fluids (plasma, urine, and amniotic fluid) for 611 pig fetuses from four different genotypes and at two different gestational stages. Note that a limit of the technique is that it is not suited to quantify lipids that form a heap of peaks in NMR spectra. In addition, a low number of metabolites related to the glycolysis and lipid pathways were found in our study because most of them were not in **ASICS** reference library. Hence, the “glycolysis pathway” and the lipid related metabolisms, which are known to be important in late gestation, were not found in our study. Despite this limitation, for the three fluids, major differences were found between the two gestational stages indicating a dramatic change in fetal metabolism between 90 and 110 days of gestation. Such a metabolic switch has also been recently reported in the metabolome of the amniotic fluid in humans between the second and the third trimesters of gestation[41]. The metabolic switch observed here is also consistent with our previous findings obtained with the same experimental design, which highlighted important variations in the muscle and intestinal transcriptomes and in the muscle and adipose tissue proteomes for a subset of the fetuses used in the present article[22, 55, 56, 66].

This study highlighted metabolomic pathways involved in the regulation of carbohydrates, amino acids, and glutathione metabolisms, which were found enriched in influential or differential metabolites. Many of the metabolites that we identified are directly related to cellular energy levels and metabolism. It is indeed critical that carbohydrate metabolism is efficient at birth to provide the newborn piglet the energy necessary to overcome hypothermia due to birth, and then the energy to meet requirements for maintenance, thermoregulation and growth[16, 30]. In addition, the different pathways related to carbohydrate metabolisms were shown to be altered during fetal development in neonates with IUGR[39]. In our study, the “galactose metabolism” was the only pathway enriched in influential or differential metabolites whatever the fluid or the statistical method. This pathway is essential for mammalian species, especially in fetal and neonatal development, because it has an important role in energy delivery[13]. All the seven metabolites (galactitol, glucose, glucose-6-phosphate, lactose, mannose, sorbitol and myo-inositol) included in the “galactose metabolism” were identified by **ASICS** in the three fluids.

Among the metabolites identified in the “galactose metabolism” pathway, the myo-inositol concentration had already been proposed as a marker of development of obesity and type 2 diabetes in human adults[14, 26] and also as a marker of IUGR in humans[1, 15] and pigs[40]. In these previous studies, a higher plasma or urine concentration of myo-inositol was associated with a higher risk of IUGR and, thus, with a lower maturity. More precisely, in pigs, NMR metabolomic profiling performed on fetuses at 110 days of gestation has demonstrated that low-weight fetuses had a higher plasma concentrations of myo-inositol than high-weight fetuses[40]. Also, Dessì and Fanos[15] have suggested that, in fetus with IUGR, greater plasma concentrations of myo-inositol may reflect an altered glucose metabolism and that they were associated with a decrease in lipid synthesis and cell proliferation through the reduction in insulin secretion. Such effects then result in a lower birth weight. Consistently, we observed lower plasma concentrations of myo-inositol in MS fetuses, considered as more mature, at both 90 and 110 dg, despite a lower concentration of myo-inositol in urine at 90 dg compared to 110 dg (no genotype effect in this fluid). This finding suggests that a more efficient glucose metabolism may be part of the greater maturity of MS piglets compared to LW piglets.

Another metabolite, the glucose, is essential to provide energy for fetal growth and development. Pig plasma concentration of glucose had already been shown to be lower in IUGR than in non-IUGR newborns[35, 51]. In our study, glucose was indeed found differential and more concentrated in MS than in LW, both at 90 and 110 dg. Therefore, both myo-inositol (less concentrated in MS) and glucose (more concentrated in MS) are possible biomarkers of maturity in plasma. Moreover, the concentrations of these two metabolites were more influenced by the paternal genotype, which is consistent with a parental imprinting mechanism[6, 43]. The role of genes, such as *IGF2* under parental imprinting during gestation has been previously described[21] and its role in the fetal glycogen synthesis has also been demonstrated[34]. However, the parental imprinting phenomenon had never been studied with metabolomic data until the current study in which we exhibit some metabolites (like myo-inositol and glucose) more concentrated in function of the paternal or maternal genotype in the reciprocal fetuses.

In addition, during the last month of gestation, glucose is stored in fetal tissues, especially in muscle and liver, in its polymerized form called glycogen. The storage of glycogen just before birth is known since Claude Bernard discovered this molecule in 1857 (reviewed by Young in 1957[67]). He showed how muscle and liver glycogen contents dramatically drop after birth in mammalian by comparing glycogen contents at birth and at a few hours after birth. In our study, glycogen was detected in urine, amniotic fluid, and to a lesser extent, in plasma and its concentration was significantly greater at 110 dg than at 90 dg in the three fluids. At birth, the piglets mainly rely on glycogen as an energy-yielding substrate before colostrum consumption[38, 54]. Studying pig fetuses close to term in relation to their value for survival at birth, Leenhouwers *et al.*[31] and Voillet *et al.*[55] had shown that glycogen content in liver and muscle increased with chance of survival. Since glycogen is a multibranched polysaccharide of glucose described as reserve in tissues, it was unexpected to find it in the fluids that we studied (*i.e.*, plasma, urine, and amniotic fluid). A possible explanation is that, as glycogen synthesis in tissues is intense, some polymers of glucose may have been released in fluids just before birth.

As for carbohydrates, many amino acids were highlighted in our analyses. Nine amino acid metabolism pathways were found enriched in influential and differential metabolites at the end of gestation: “alanine, aspartate and glutamate metabolism”, “aminoacyl tRNA biosynthesis”, “arginine and proline metabolism”, “cyanoamino acid metabolism”, “cysteine and methionine metabolism”, “glycine, serine and threonine metabolism”, “lysine biosynthesis”, “lysine degradation” and “valine, leucine and isoleucine biosynthesis”. They all respond to a necessity for fetal development and maturation since amino acids have nutritional, physiological and regulatory roles. Twenty amino acids are involved in these pathways[47]. In our study, these pathways were enriched with 15 differential amino acids, including 5 essential amino acids (*i.e.*, amino acids that cannot be synthetized by animals) and 10 non essential amino acids (*i.e.*, that can be synthetized by animals). Amino acids of the “arginine and proline metabolism” (arginine, asparagine, aspartate, glutamate, glutamine, ornithine, and proline) have already been well studied in a gestational context because of their essential role in growth and development of the fetus both in humans and pigs[36].

In early pregnancy, arginine concentration in the amniotic fluid has been described as positively correlated with birth weight, length and head circumference of babies[7]. Moreover, Wu *et al.*[61] showed that an arginine supplementation of sows during the gestation decreases the stillbirth rate and the IUGR risk. These two previous studies are in favor of an important role of arginine fetal maturation. They are also consistent with our findings: arginine was identified by mixed models as differential. It was found more concentrated at 110 dg than at 90 dg in all fluid, even though an earlier study[35] had shown a decrease in plasma concentration of arginine between 90 and 110 dg.

As for arginine, glutamine is also highly concentrated in amniotic fluid mainly in early gestation[62]. At the end of gestation, the amniotic fluid serves as a nutritional reservoir for the fetus and, thus, the uptake of glutamine by the fetus may induce a decrease in glutamine concentration in the amniotic fluid[36]. Hence, glutamine concentration is usually considered as a limiting factor of fetal growth and a lower concentration is known to be associated with higher IUGR risk. This was confirmed by our study in which glutamine was not found at 110 dg in amniotic fluid. Proline, which is also in the “arginine and proline metabolism” pathway, is less frequently used in sow nutrition than arginine and glutamine[37]. However, its important role in polyamine synthesis during the pig gestation has previously been demonstrated[60]. As expected, the higher concentration of proline in MS compared to LW at 90 and 110 dg in plasma could be related to the lower maturity of LW piglet and to a potential delay in development already identified in those fetuses[22, 55, 56, 66]. In addition, in plasma, crossbred fetuses have intermediate proline concentrations, with no specific maternal or paternal effect.

Serine, glycine and guanidinoacetate are metabolites involved in the “one-carbon metabolism” (this metabolism is not a KEGG pathway so was not studied for enrichment). This metabolism participates to the DNA methylation by providing methyl groups[36]. As for imprinting genes, DNA methylation is an important epigenetic mechanism of fetal gestation. Previous studies have shown the association between IUGR and epigenetic alterations[36]. Lin *et al.*[35] have also shown that the concentration of serine significantly decreases between 90 dg and 110 dg in pigs. Moreover, a higher concentration of serine and glycine in plasma has been reported in IUGR rat fetuses compared to normal weight fetuses[42]. This is consistent with our findings: glycine and serine concentrations were found differential and with a higher concentration at 90 dg than at 110 dg in plasma. Glycine concentration was also found more concentrated in LW than in MS at both 90 and 110 dg. Moreover, guanidinoacetate exhibit a maternal effect: concentrations of this metabolite in fetuses with a LW mother are higher than in fetuses with a MS mother whatever the stage of gestation. For glycine, the same maternal effect is observed at 90 dg. These metabolites are precursors of the creatine (see Fig. 3), which is known to be involved in energy metabolism and development of skeletal muscles[8, 58]. In plasma, both creatine and creatinine were found differential and were more concentrated in MS than in LW at 110 dg. Other studies on IUGR had reported a higher concentration for creatine and creatinine in IUGR fetal pigs[35] or newborns[14], in contrast to our findings. However, in our study, it can be noted that the concentration of creatinine in plasma changes differently according to the genotype. Indeed, the concentration was approximately the same, whatever the genotype, at 90 dg but, for LW, the concentration sharply decreased and, for MS, it sharply increased at 110 dg compared to 90 dg.

Furthermore, as the production of oxidants increases during the gestation due to the cell proliferation, it is necessary to have an efficient “glutathione metabolism” that plays a role in oxidative defense[63, 64]. An increase in oxidative stress has already been associated with IUGR or preterm infants[27–29, 63]. As expected, in our study, the “glutathione metabolism” was found enriched in differential metabolites in urine. Reduced glutathione is formed from glutamate, cysteine and glycine and protects cells against oxidative damage by removing hydrogen peroxide[47]. More precisely, the oxidized glutathione was detected only in MS and found more concentrated at 90 dg than at 110 dg. On the contrary, the reduced glutathione was found more concentrated at 110 dg than at 90 dg in MS and was almost not detected in LW. Altogether, these results suggest a better oxidative defense of MS compared to LW. Interestingly, the concentration of pyroglutamate, another metabolite involved in the “glutathione metabolism”, had already been shown more concentrated in the plasma of IUGR fetuses compared with the normal birth weight group, a fact that was likely due to a reduced synthesis of glutathione[59]. In addition, the pyroglutamate has been considered as a biomarker for IUGR in fetal plasma[35]. However, this is not a clearly established fact. For instance, Jackson *et al.*[29] have shown that the pyroglutamate/creatinine ratio in urinary excretion just after birth is greater for preterm infants. We observed the same trend in the late gestation: the pyroglutamate was found more concentrated at 110 dg than at 90 dg in plasma and the pyroglutamate/creatinine ratio was higher in LW than in MS in urine.

## Conclusions

As pointed by Leite and Cecatti[33], one of the main interest of metabolomic approaches is to identify criteria of an optimal growth for fetuses and newborns, that can go beyond using only the birth weight. This should allow to discriminate between fetuses with really restricted growths and small weight newborns (that are just “constitutionally small” but with good further outcomes). In particular, proline and myo-inositol are two promising metabolites for the characterization of piglet maturity. They illustrate the importance of the amino acid and carbohydrate metabolisms for the fetal development in late gestation.

## Methods

### Ethics Statement

All fluids from pig fetuses during the PORCINET project (ANR-09-GENM-005-01, 2010–2015). The experiment authorization number of the experimental farm GenESI (Pig phenotyping and Innovative breeding facility, doi:10.15454/1.5572415481185847E12) is A-17-661. The procedures and the animal management complied with European Union legislation (Directive 2010/63/EU) and the French legislation in the Midi-Pyrénées Region (Decree 2001-464). The ethical committee of the Midi-Pyrénées Regional Council approved the experimental design (authorization MP/01/01/01/11).

### Animals and plasma, urine and amniotic liquid sampling

Plasma, urine and amniotic fluid samples were obtained from 611 pig fetuses at two gestational stages (90 and 110 days, average gestation term at 114 days). MS and LW sows were inseminated with mixed semen (LW and MS) so that most litters were composed of purebred fetuses (LW or MS) and crossbred fetuses (LW×MS from MS sows and MS×LW from LW sows). MS and LW breeds were chosen as two extreme breeds for piglet mortality at birth, a better survival rate being observed in MS piglets. The detailed experimental design was previously described in Voillet *et al.*[56] and is summarized in Supplementary Fig. S5. 329 fetuses had a LW mother and 282 fetuses have a MS mother. The fetuses were obtained by caesarean section. Fetuses were weighed (statistics on weights are provided in Table 3), showing an average weight larger for LW than for MS despite their lower maturity.

**Table 3:** Fetus weights at 90 and 110 days of gestation by genotype (mean *pm* standard deviation in grams).

| Genotype | 90 days of gestation | 110 days of gestation |
| --- | --- | --- |
| LW | 619 ± 141 | 1171 ± 323 |
| MS×LW | 633 ± 91 | 1292 ± 197 |
| LW×MS | 579 ± 114 | 1092 ± 201 |
| MS | 490 ± 86 | 910 ± 101 |

After laparotomy of the sow, blood (approximately 5 mL) was collected individually on piglets from the umbilical artery via a 21-gauge needle and a 5 mL syringe and placed in heparinized tubes. After section of the umbilical cord, the fetus was euthanized [66]. Plasma was prepared by low-speed centrifugation (2,000 g for 10 min at 4°C) and stored at −80°C until further analysis. The amniotic fluid (10 mL) was collected during the caesarean and immediately centrifuged (2,000 g for 10 min at 4°C) to discard cell debris and stored at −20°C until further analysis. The urine samples were collected during the fetuses dissection directly in the bladder with a 5 mL syringe, immediately frozen to avoid contamination and stored at −80°C until further analysis.

### Nuclear magnetic resonance

Detailed protocol of sample preparation, spectra acquisition and preprocessing can be found in Lefort *et al.*[32]. In summary, each sample of plasma and amniotic fluid (200 *μ*L) was diluted in 500 *μ*L deuterated water (D_2_O) and centrifuged without the addition of internal standard to improve spectra quality. For urine, 200 *μ*L of phosphate buffer prepared in deuterated water (0.2 M, pH 7.0) were added to 500 *μ*L of urine, vortexed, centrifuged at 5000 g for 15 min, and 600 *μ*L transferred to 5 mm NMR tube. All ^1^H NMR spectra were acquired on a Bruker Avance DRX-600 spectrometer (Bruker SA, Wissembourg, France) operating at 600.13 MHz for ^1^H resonance frequency and at 300K using the Carr-Purcell-Meiboom-Gill (CPMG) spin-echo pulse sequence. The Fourier transformation was calculated on 64,000 points. Moreover, all spectra were phased, baseline corrected and then calibrated on the resonance of lactate (1.33 ppm) using Topspin (V2.1, Bruker, Biospin, Munich, Germany). The regions corresponding to water resonance (5.1–4.5 ppm) and urine (6.5–6.0 ppm) were excluded to eliminate artifacts of residual water and urine.

### Metabolite identification and quantification

Before quantification, spectra were normalized by the area under the curve and aligned with pre-processing functions available in the Bioconductor R package **ASICS**[32] (version 2.0.0). Then, the metabolites in all fluids were identified and quantified using the ASICS method available in the same package. The quantification was performed using the default reference library provided in the package and was processed independently for every fluid but with the same maximum chemical shift allowed, which was set to 0.02. Moreover, library alignment was improved (as compared to what is described in Lefort *et al.*[32]) by using a global quality control criterion: the correlation between quantifications and targeted buckets of the spectra was maximized to choose the best alignment between all peaks. Finally, identified metabolites were kept only for the metabolites that had at least 25% of quantifications larger than 0 in at least one condition (stages of gestation and genotypes). The others were removed from the list of identified metabolites (quantification set to 0).

### Spectra quality control

In Lefort *et al.*[32], plasma relative quantifications had already been validated using biochemical targeted dosages of three metabolites (glucose, fructose and lactate) in a subset of the samples. Moreover, results of a an Orthogonal Projections to Latent Structures Discriminant Analysis[52] (OPLS-DA) based on the classical bucket approach were compared with those of an OPLS-DA based on metabolite quantifications. The comparison showed a good reproducibility and a similar discriminative power between conditions for the bucket and the quantification approaches, ensuring a minimal loss of information resulting from the quantification preprocessing.

Principal Component Analyses (PCA) were used to detect potential outliers and batch effects due to experimental covariates: sex, breeding batch, and sow. All plots are available in Supplementary Fig. S6-S9. PCA did not identify any sex, batch or experimental effect but a sow effect was clearly visible and thus later included in the analyses when possible.

### Multivariate exploratory analyses

All statistical analyses were performed with R (version 3.6.0)[45]. The effect on the metabolome of the stage of gestation (*i.e.*, 90 dg and 110 dg) was first investigated with OPLS-DA[52]. Three OPLS-DA were thus performed independently on each fluid to find the metabolites with the highest discriminant power between the two gestational stages: the most influential metabolites, *i.e.*, the metabolites with a VIP index superior to 1, were extracted. The relevance of the results was ensured by estimating the predictive power of each model with a 10-fold cross-validation error.

### Univariate differential analyses with mixed models

OPLS-DA is limited to the study of one factor with only two levels so we complemented it with more complete analyses based on mixed models, which are able to incorporate multiple effects, including the random effect of the sow used as a proxy for the effect of the uterine environment. This effect must not be mistaken with a parental effect coming from the genotype. Mixed models were used to find metabolites with differential concentrations between conditions (gestational stages and genotypes), by fitting the following model for each fluid and each metabolite:

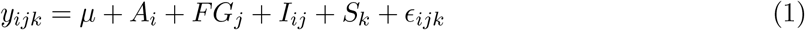

with *y*_*ijk*_ the vector of metabolite concentrations for gestational stage *i* (*i* ∈ {d90, d110}), genotype *j* (*j* ∈ {LW, LW × MS, MS × LW, MS}) and mother (sow) *k*. In this model, *μ* is the mean effect, *A*_*i*_ the fixed effect of the gestational stage, *FG*_*j*_ is the fixed effect of the genotype, *I*_*ij*_ is the effect of the interaction between the gestational stage and the genotype, 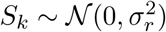 is the random effect of the sow and 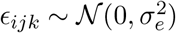 is a noise term.

For all metabolites, this model was tested against the model with only the sow effect (*y*_*ijk*_ = *μ* + *S*_*k*_ + *ϵ*_*ijk*_) with a Fisher test. *p*-values were then adjusted with the Benjamini and Hochberg (FDR) correction [5]. Finally, using the same methodology as Voillet *et al.* [56], each differential metabolite (*i.e.*, the metabolites with an adjusted *p*-value smaller than 0.05) was associated with one of the following sub-model:

- complete: *y*_*ijk*_ = *μ* + *A*_*i*_ + *FG*_*j*_ + *I*_*ij*_ + *S*_*k*_ + *ϵ*_*ijk*_
- additive: *y*_*ijk*_ = *μ* + *A*_*i*_ + *FG*_*j*_ + *S*_*k*_ + *ϵ*_*ijk*_
- only stage: *y*_*ijk*_ = *μ* + *A*_*i*_ + *S*_*k*_ + *ϵ*_*ijk*_
- only genotype: *y*_*ijk*_ = *μ* + *FG*_*j*_ + *S*_*k*_ + *ϵ*_*ijk*_

Contrary to the approach that would have consisted in testing each effect of the complete model (stage effect, genotype effect and interaction) independently, selecting the best fit sub-model avoids overfitting and leads to obtain the best set of relevant effects for every metabolite. Since the four models described above are not nested (only stage and only genotype models are not), this selection cannot be performed using a standard Fisher test so we used a model selection approach instead and selected the model with the minimum Bayesian Information Criterion (BIC)[49].

### Pathway enrichment

Pathway enrichment analysis was performed with the web-based tool suite MetaboAnalyst[12] (version 4.0) more precisely with the MetPA module[65]. *Sus scrofa* pathways were not available thus the *Homo sapiens* KEGG pathways were used instead, as a reference. We also checked with KEGG database the difference between human and pig pathways which were found to be almost identical. This confirmed the relevance of using the human pathways in MetaboAnalyst. Moreover, hypergeometric test and *p*-value correction based on FDR were used to extract pathways enriched into influential or differential metabolites. This analysis was carried out independently for each fluid on OPLS-DA and mixed model results.

## Acknowledgements

This project received financial support from French National Agency of Research (PORCINET grant ANR-09-GENM005). The PhD fellowship of Gaelle Lefort is supported by the Digital Agriculture Convergence Lab (#DigitAg, http://www.hdigitag.fr/, ANR-16-CONV-0004), by INRAE Mathematics and Computer Science Division, by INRAE Animal Genetics Division and by INRAE Animal Health Divison. The funders had no role in the study design, analyses, results interpretation and decision to publish. The authors are very grateful to the staff of experimental pig facilities (INRA, 2018. Pig phenotyping and Innovative breeding facility, doi: 10.15454/1.5572415481185847E12) and Marie-Christine Père (PEGASE, INRAE) for expert assistance in surgeries. They also thank all the participants from GenPhySE, PEGASE and GenESI laboratories (INRAE) for sample and data collection. The authors also thank Anne Poujol for her veterinarian assistance.

## Author contributions statement

L.L., H.Q. and L.C. conceived and designed the study. L.L. supervised the PORCINET project. Y.B. supervised the performance testing, from animal production to biological sampling. N.I. was the project data manager. C.C. and A.P. managed and performed the metabolomic experiments. G.L., R.S. and N.V. developed the statistical models and performed the data analyses. G.L., R.S., H.Q., N.V. and L.L. wrote the article. All authors reviewed and approved the manuscript.

## Data avaibility

The data supporting the results of this article are available in the MetaboLights database [**?**]: MTBLS1541 (www.ebi.ac.uk/metabolights/MTBLS1541).

## Additional information

**Correspondence** and requests for materials should be addressed to G.L. and L.L.

