## Supplementary Material for "The maturity in fetal pigs using a multi-fluid metabolomic approach"

### Supplementary material of the article "The maturity in fetal pigs using a multi-fluid metabolomic approach"

G. Lefort *et al.*

#### Contents

**S1 Supplementary tables** **2**

**S2 Supplementary figures** **4**

### S1 Supplementary tables

Table S1. Metabolites identified in each fluid.

| metabolite names in ASICS | alternative names | plasma | urine | amniotic fluid |
|---|---|---|---|---|
| 1,3-Diaminopropane | | X | X | |
| 2-AminoAdipicAcid | 2-Aminoadipate | X | X | X |
| 2-AminobutyricAcid | | X | | |
| 2-Deoxycytidine | | | X | X |
| 2-Oxoglutarate | | X | | X |
| 2-Oxoisovalerate | | X | | |
| 3-Hydroxybutyrate | | | X | |
| 3-Methyl-L-Histidine | | | X | X |
| 3-MethyladipicAcid | | | X | |
| 4-AminoHippuricAcid | | X | | |
| 4-HydroxyphenylAceticAcid | | X | | |
| 5-AminoValericAcid | 5-Aminopentanoate | | X | X |
| alpha-HydroxyisobutyricAcid | | | | X |
| ArgininosuccinicAcid | Arginosuccinate | | X | |
| AscorbicAcid | Ascorbate | | | X |
| Azelaic Acid | | | | X |
| Betaine | | X | X | X |
| Cadaverine | | X | X | X |
| CholineChloride | Choline | X | X | |
| Citrate | | | | X |
| Creatine | | X | | X |
| Creatinine | | X | X | X |
| D-Fructose | Fructose | X | X | X |
| D-Fucose | | X | X | X |
| D-Galactose | | | | X |
| D-GluconicAcid | Gluconate | X | X | X |
| D-Glucose | Glucose | X | X | X |
| D-Glucose-6-Phosphate | Glucose-6-phosphate | X | X | X |
| D-GlucuronicAcid | Glucuronate | X | X | X |
| D-Maltose | Maltose | | X | |
| D-Mannose | Mannose | X | X | X |
| D-Sorbitol | Sorbitol | X | X | X |
| DehydroAscorbicAcid | | X | X | X |
| Ethanolamine | | X | X | X |
| EthylmalonicAcid | | X | | |
| Galactitol | | X | X | X |
| GlycericAcid | Glycerate | | X | X |
| Glycerol | | X | X | X |
| Glycerophosphocholine | | X | | X |
| Glycogen | | X | X | X |
| GuanidinoaceticAcid | Guanidinoacetate | X | X | X |
| Hypotaurine | | | | X |
| IsocitricAcid | | X | X | X |
| IsovalericAcid | | X | | |
| L-Alanine | Alanine | X | X | X |
| L-Arabitol | | X | X | X |
| L-Arginine | Arginine | X | X | X |
| L-Asparagine | Asparagine | | X | |
| L-Aspartate | Aspartate | X | X | X |
| L-Carnitine | | X | | |

**Table S1 continued from previous page**

| metabolite names in ASICS | alternative names | plasma | urine | amniotic fluid |
|---|---|---|---|---|
| L-Citrulline | Citrulline | X | X | X |
| L-Cysteine | Cysteine | | | X |
| L-Cystine | | X | X | X |
| L-GlutamicAcid | Glutamate | X | X | X |
| L-Glutamine | Glutamine | | X | X |
| L-Glutathione-oxidized | Oxidized glutathione | | X | X |
| L-Glutathione-reduced | Reduced glutathione | | X | X |
| L-Glycine | Glycine | X | X | X |
| L-Isoleucine | Isoleucine | X | | X |
| L-Leucine | Leucine | X | | X |
| L-Lysine | Lysine | | X | X |
| L-Proline | Proline | X | X | X |
| L-Serine | Serine | X | X | X |
| L-Threonine | Threonine | X | X | X |
| L-Valine | Valine | X | | X |
| Lactate | | X | X | X |
| Lactose | | X | X | X |
| Levoglucosan | | | X | X |
| MandelicAcid | | X | | |
| Methanol | | | X | X |
| Methylguanidine | | | X | |
| Myo-Inositol | | X | X | X |
| N-Acetylglycine | | X | X | |
| PantothenicAcid | Pantothenate | X | | X |
| Phenethylamine | | X | | |
| PropyleneGlycol | | X | X | |
| PyroglutamicAcid | Pyroglutamate | X | X | X |
| S-Acetamidomethylcysteine | | X | X | X |
| SaccaricAcid | | | X | X |
| Sarcosine | | X | X | |
| SebacicAcid | | X | | X |
| Spermidine | | | X | X |
| Taurine | | X | | X |
| Threitol | | X | X | X |
| ThreonicAcid | Threonate | X | X | X |
| TMAO | | X | X | X |
| trans-4-Hydroxy-L-Proline | Hydroxyproline | X | | |
| UDPG | | | X | |
| Uridine | | | X | X |
| Xylitol | | X | X | X |

#### S2 Supplementary figures

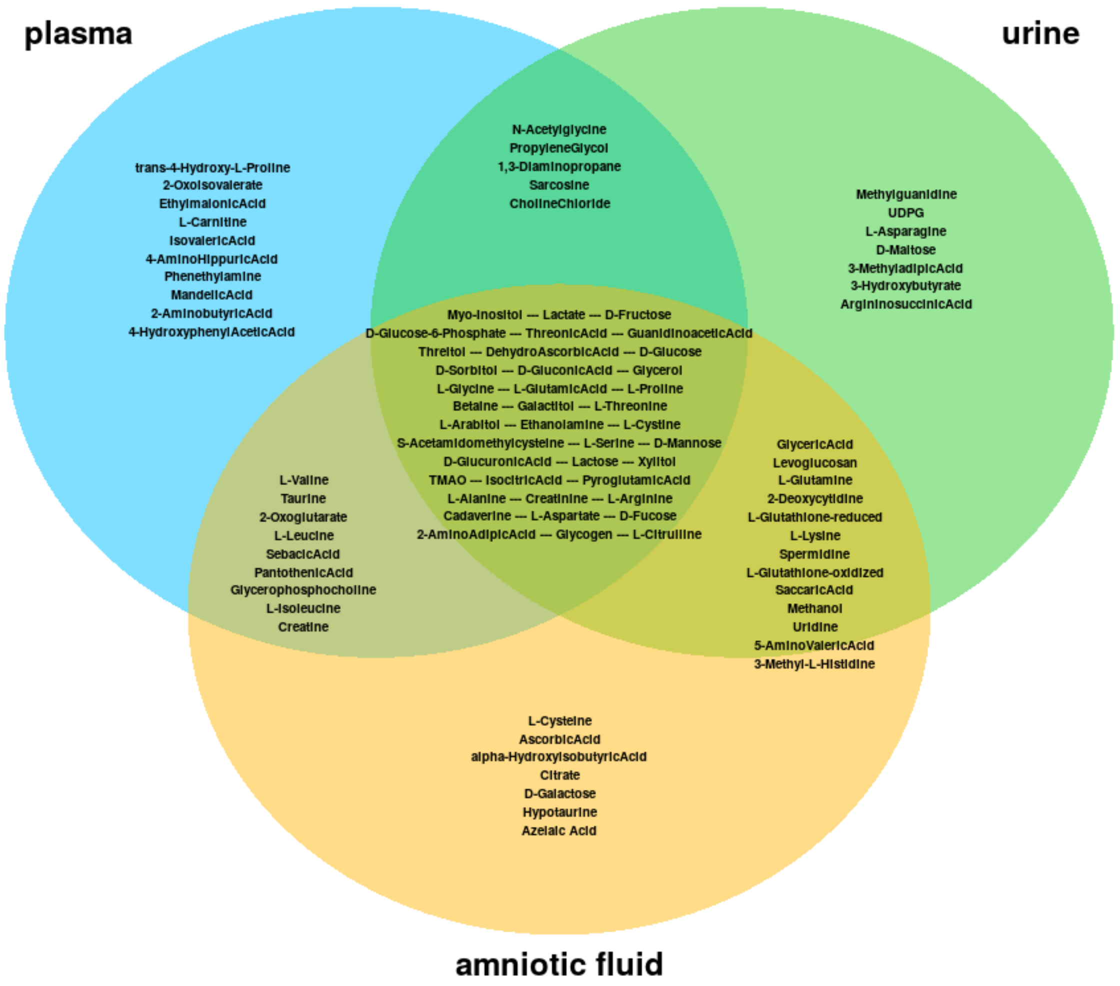


**Fig. S1.** Metabolites identified with **ASICS** package in each fluid.

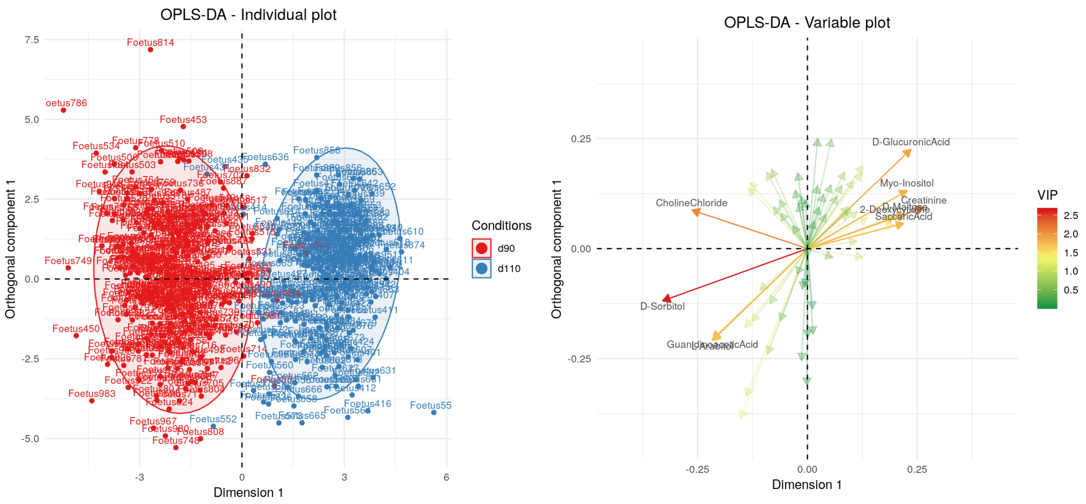


**(a)** OPLS-DA on urine spectra

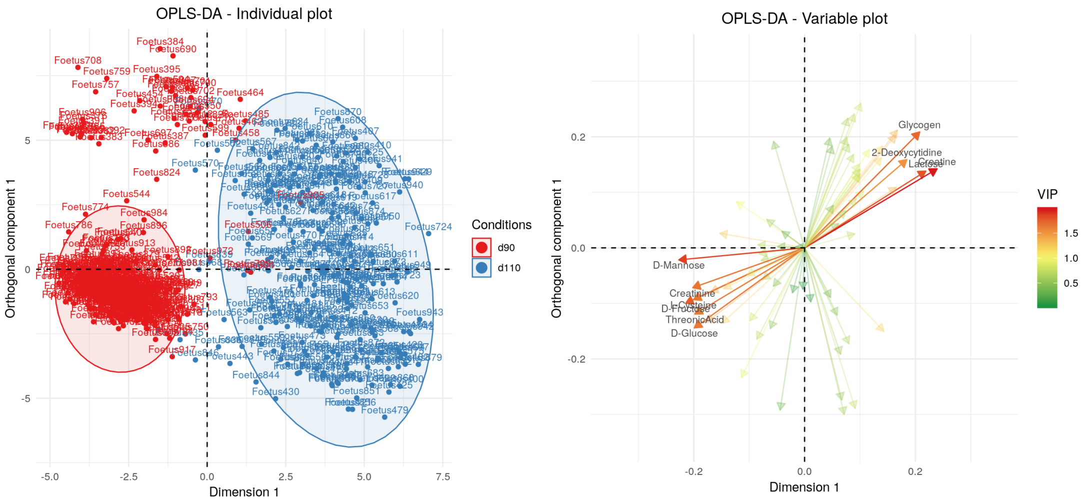


**(b)** OPLS-DA on amniotic fluid spectra

**Fig. S2.** Results of OPLS-DA.

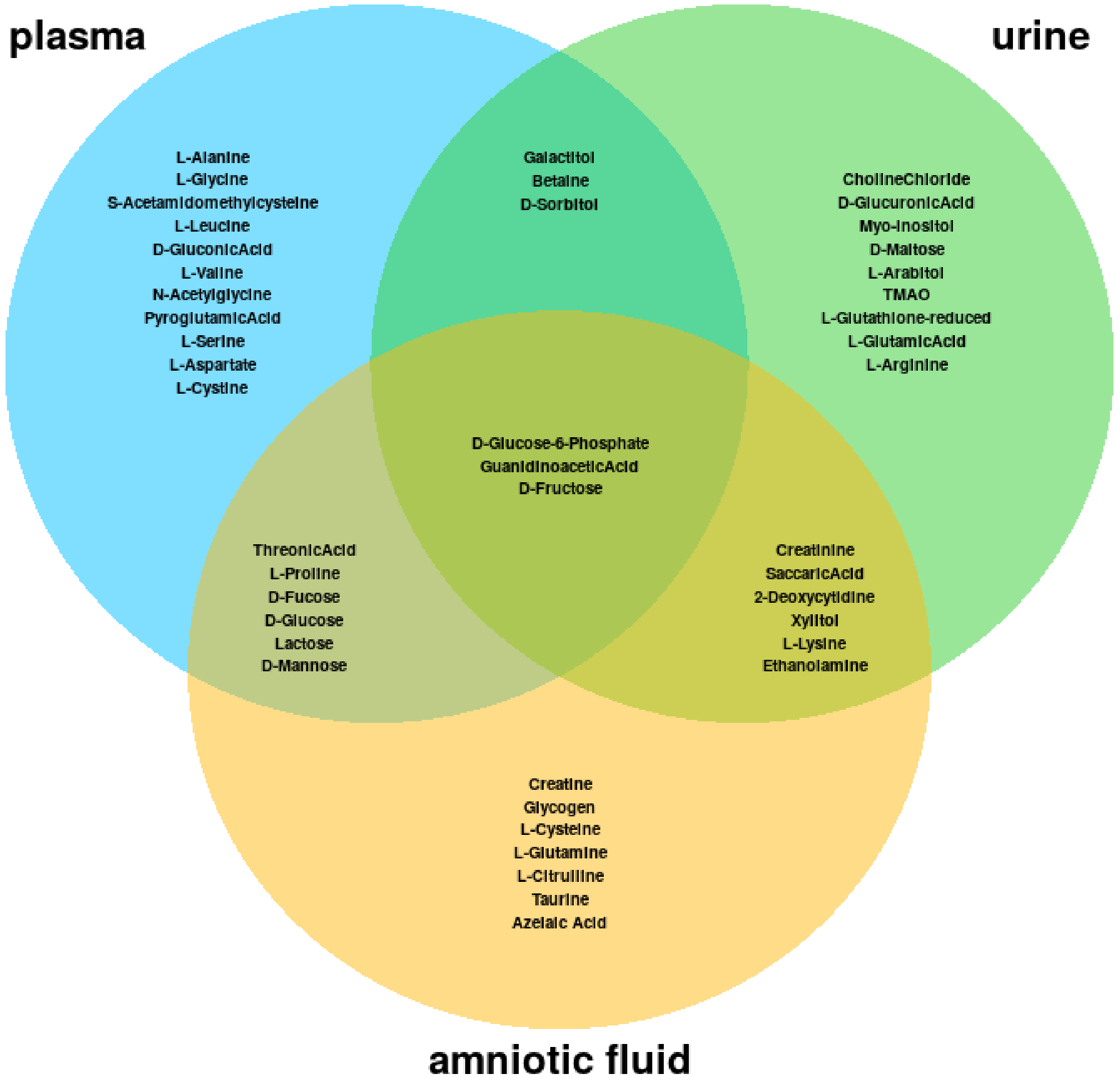


**Fig. S3.** Influential metabolites detected by OPLS-DA.

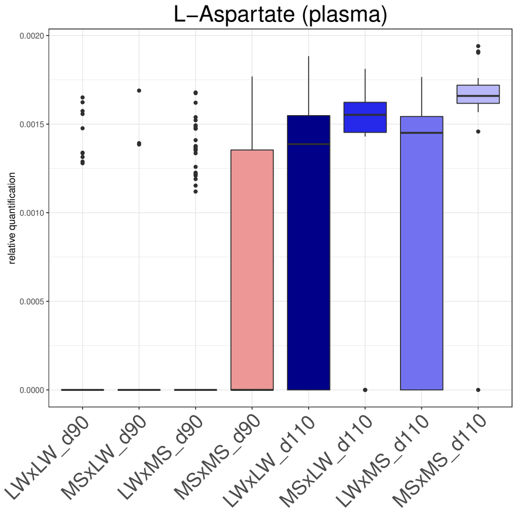


**(a)** Aspartate concentrations in plasma by gestational stage and genotype

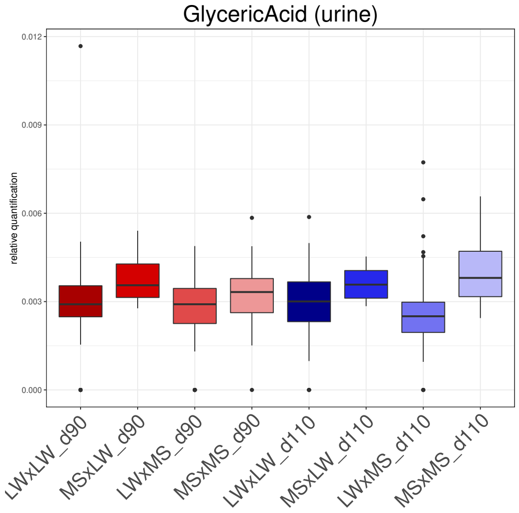


**(b)** Glycerate concentrations in urine by gestational stage and genotype

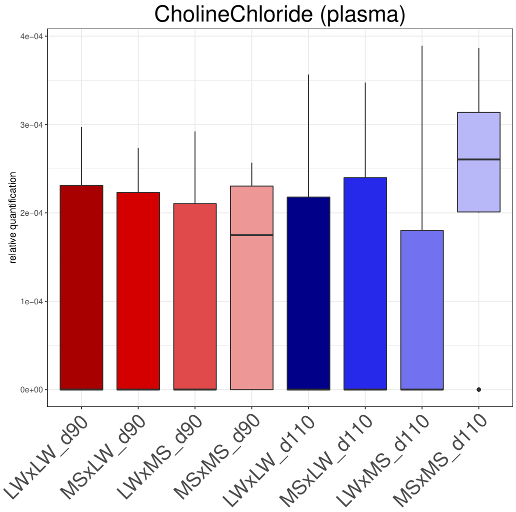


**(c)** Choline concentrations in plasma by gestational stage and genotype

**Fig. S4.** Paternal effect for aspartate in plasma and glycerate in urine and effect of the pure MS genotype for choline in plasma.

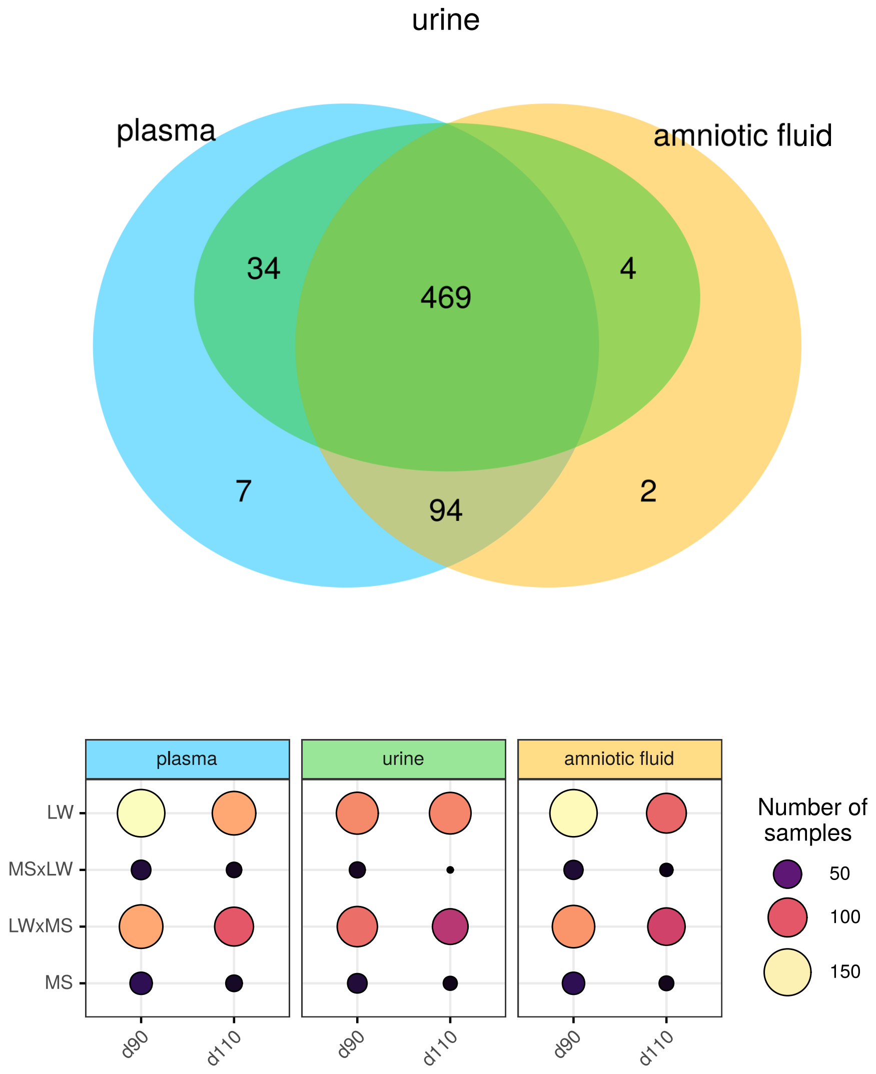


**Fig. S5.** Summary of the design experiment: number of samples by fluids, stage of gestation and genotype.

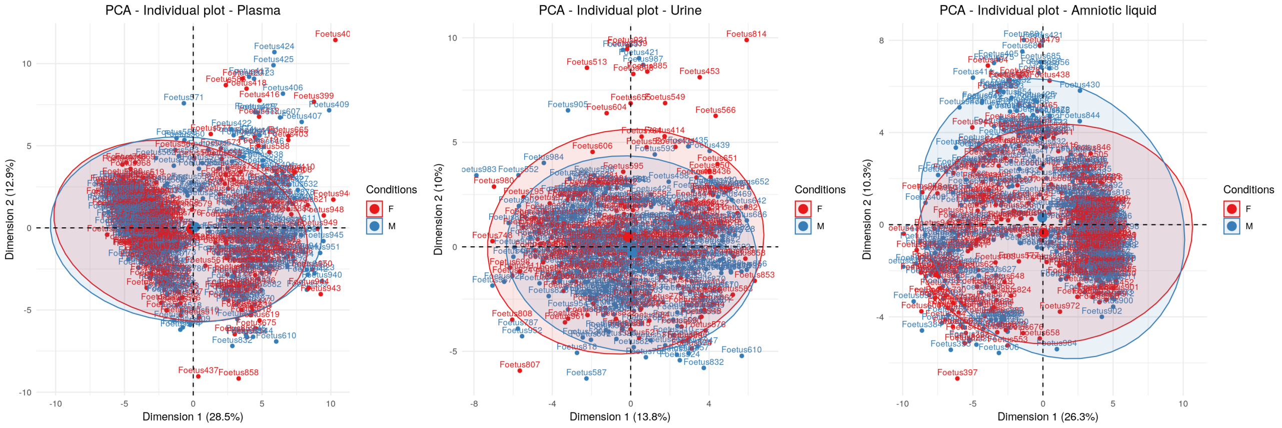


**Fig. S6.** PCA on quantifications of metabolitess colored by gender (red: females, blue: males).

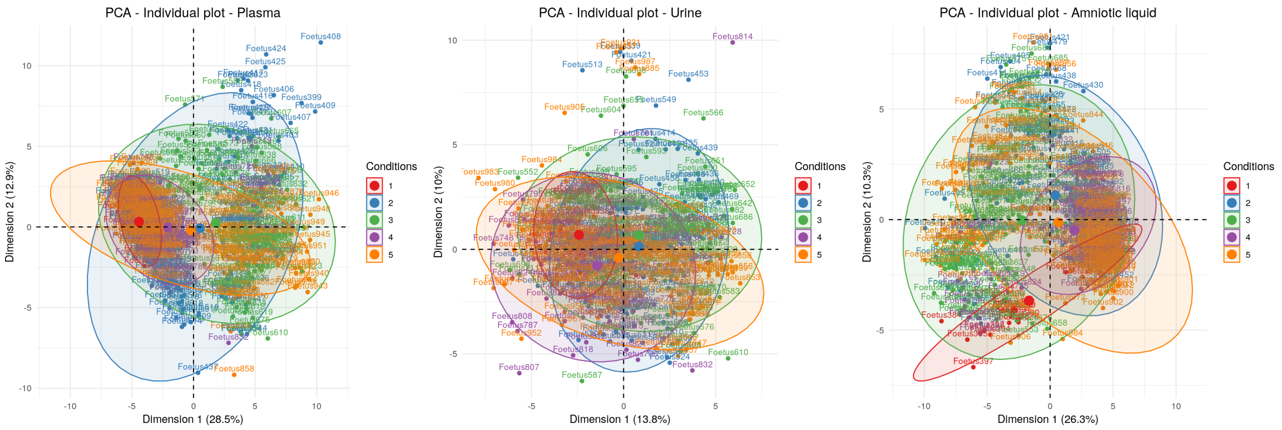


**Fig. S7.** PCA on quantifications of metabolites colored by batch effect.

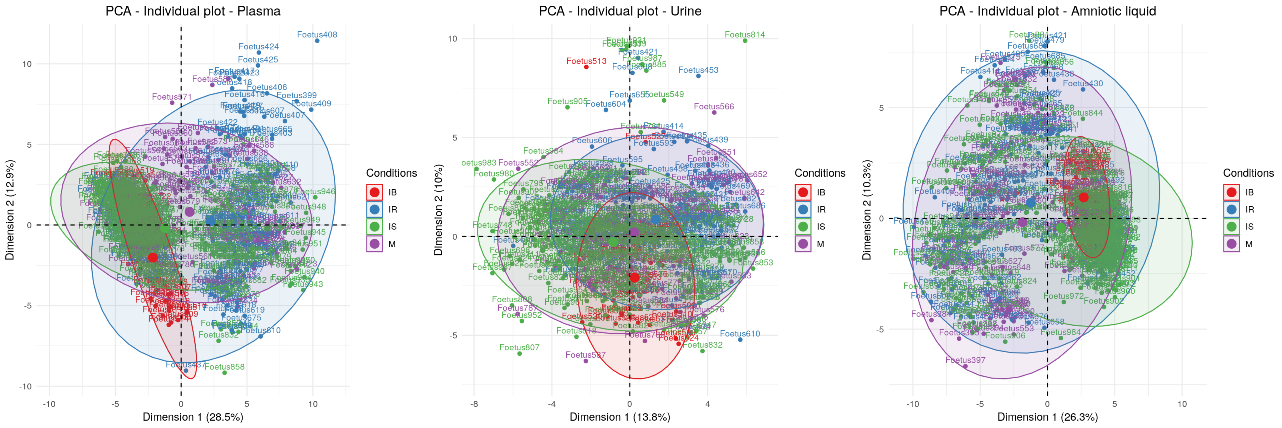


**Fig. S8.** PCA on quantifications of metabolites colored by anesthesia method effect (IB : intubated with manual ventilation, IR: intubated under respirator, IS: intubated in spontaneous breathing and M: mask).

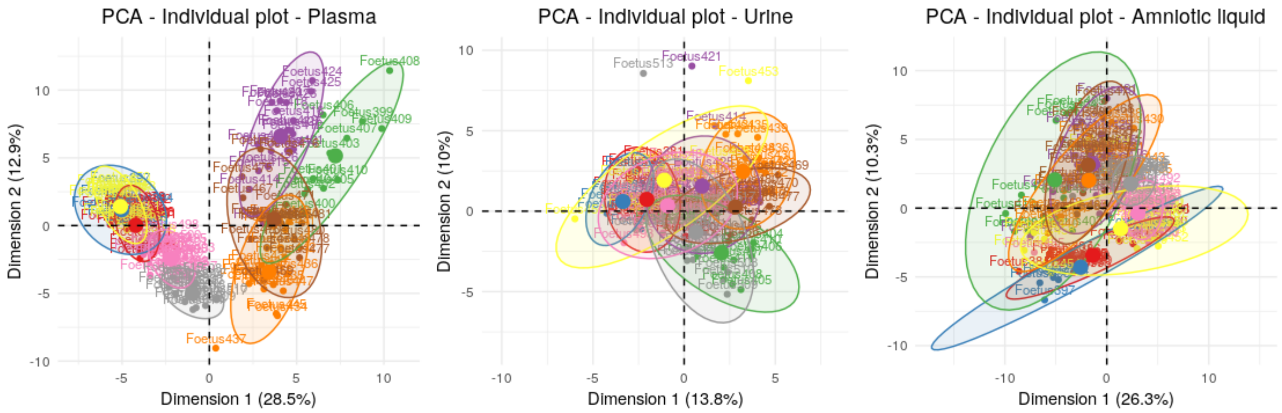


**Fig. S9.** PCA on quantifications of metabolites colored by sow. For the sake of clarity, only the first nine sows are represented.
